# Combining multiple genetic estimates of *N*_*e*_

**DOI:** 10.1101/2025.06.26.661823

**Authors:** Robin S. Waples

## Abstract

Researchers often use multiple genetic methods to estimate contemporary effective population size (*N*_*e*_), but few formally combine estimates despite potential benefits for increasing precision. Maximizing benefits requires an optimal, inverse-variance weighting scheme. Methods should be estimating the same parameter, which can be appropriate either for estimates using the same method applied to different time periods, or estimates using different methods applied to the same time period. Previous approaches focused on 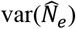 for weighting, but that is problematical because 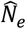 is highly skewed and can be infinitely large. A new approach is described using weights inversely proportional to 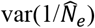, which is the drift signal that 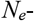 estimation methods respond to. The distribution of 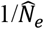 is close to normal even when 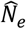 assumes extreme values. Benefits are maximized under three general conditions: estimators have approximately equal variances; they are uncorrelated or have weak positive correlations; individual estimates have low precision (i.e., if data are limited and/or true *N*_*e*_ is large). Analytical and numerical results demonstrate that: (1) existing theory allows robust estimates of 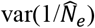 for the temporal and LD methods, which provide independent information about *N*_*e*_ – both of which facilitate optimally combining those methods; (2) estimates for the LD and sibship methods are essentially uncorrelated when data are limited but can be strongly positively correlated in genomics-scale datasets. General theory predicting 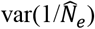 for the sibship method is lacking, but values for specific scenarios have been published. New software (*C**ombo**N**e*) is introduced to calculate combined estimates.

## 1 INTRODUCTION

Effective population size (*N*_*e*_) is one of the most important parameters in evolutionary biology, and interest in estimating *N*_*e*_ has grown rapidly in recent years (Palstra and Fraser 2012; Wang et al. 2016; Clarke et al. 2024; Waples 2025). This upward trend is likely to continue into the foreseeable future, given that (1) parties to the Convention on Biological Diversity recently committed to tracking genetic diversity for all species, and (2) effective size was identified as a key metric for monitoring within-population genetic diversity (CBD 2022; Mastretta-Yanes et al. 2024). A wide range of genetic methods for estimating *N*_*e*_ have been proposed, and collectively the time frames considered by these methods extend from the present to thousands of generations in the past (reviewed by Nadachowska-Brzyska et al. 2022). For applied conservation and management, the most relevant estimates are those for contemporary *N*_*e*_, which roughly corresponds to the time frame encompassed by the samples. The three most widely-used methods to estimate contemporary *N*_*e*_ are the temporal method (which measures changes in allele frequency between two or more samples spaced in time) and two methods that require only a single sample: the sibship method of Wang (2009), and a bias-corrected version of the method based on linkage disequilibrium (LD; Waples and Do 2008). Two other single-sample methods—the heterozygote-excess method of Pudovkin et al. (1996) and the molecular coancestry method of Nomura (2008)—have not performed well in performance evaluations (Do et al. 2014; Wang 2016) and are not widely used.

Precision is often a limiting factor for genetic methods to estimate contemporary *N*_*e*_, especially when true *N*_*e*_ is large. Furthermore, many authors report *N*_*e*_ estimates from more than one method. This suggests that formally combining information from multiple methods could lead to more robust overall estimates. To illustrate potential benefits, consider two fundamental properties of a variance. First, the variance of the product of a random variable (*X*) and a constant (*a*) is

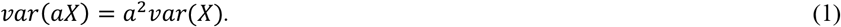

Second, the variance of the sum of two random variables is

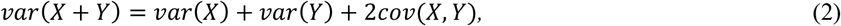

where *cov*(*X, Y*) is the covariance of *X* and *Y*. Combining Equations 1 and 2, we have

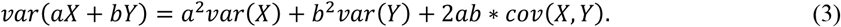

Now assume that both *X* and *Y* are unbiased estimators of the same parameter *θ*, and let *a* and *b* be standardized such that *a*+*b*=1. Then *Z* = *aX* + *bY* is a weighted average of *X* and *Y*, with *a* and *b* being relative weights. Consider a simple example where *var*(*X*) = *var*(*Y*), so *X* and *Y* are weighted equally (*a*=*b*=0.5). Substituting into Equation 3:

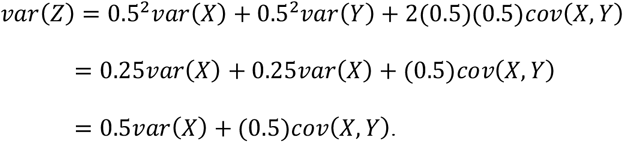

If *X* and *Y* are independent (hence *cov*(*X, Y*) = 0), then *var*(*Z*) = 0.5*var*(*X*); that is, the variance of the weighted mean would be half that of either individual estimator. More generally, Cochrane (1937; see also Shahar 2017) showed that if *X* and *Y* are independent and normally distributed, and if the weights are chosen to be inversely proportional to variances, such that

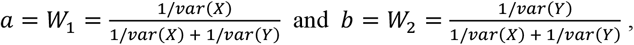

then *Z* = *W*_1_*X* + *W*_2_*Y* is an unbiased estimator of *θ* and *var*(*Z*) = *var*(*X*)*var*(*Y*)/[*var*(*X*) + *var*(*Y*)], which is less than or equal to the smaller of *var*(*X*) and *var*(*Y*).

Waples and Do (2010) showed how to combine estimates from the LD and temporal methods, but combined estimates are rarely presented, even by authors who report results from more than one method. One difficulty is that Waples and Do (2010) calculated weights based on published formulas for 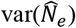 for the temporal method (by Pollak 1983) and for the LD method (by Hill 1981). However, the distribution of 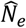 can be highly skewed and the estimates can take infinitely large or even negative values, which makes calculating 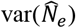 problematical except in some scenarios where the distribution of 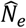 is quasi-normal.

The goal here is to develop a more robust way for combining multiple estimates of effective size, with the weights being inversely proportional to 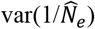 rather than 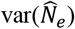. This simple change has several advantages compared to the approach outlined by Waples and Do (2010). First, 1/*N*_*e*_ is the drift signal that all genetic methods to estimate contemporary effective size are sensitive to (Wang 2001, 2009). Second, 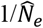 is approximately normally distributed even when 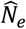 decidedly is not (Figure 1), which facilitates setting confidence intervals (CIs). Third, 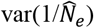has a robust analytical solution for both the temporal and LD methods, as a function of parameters that are known (sample sizes of loci and individuals; elapsed time between samples) or can be estimated (true *N*_*e*_). Here I consider how to combine multiple estimates of effective size, both within and between methods.

**Figure 1.**
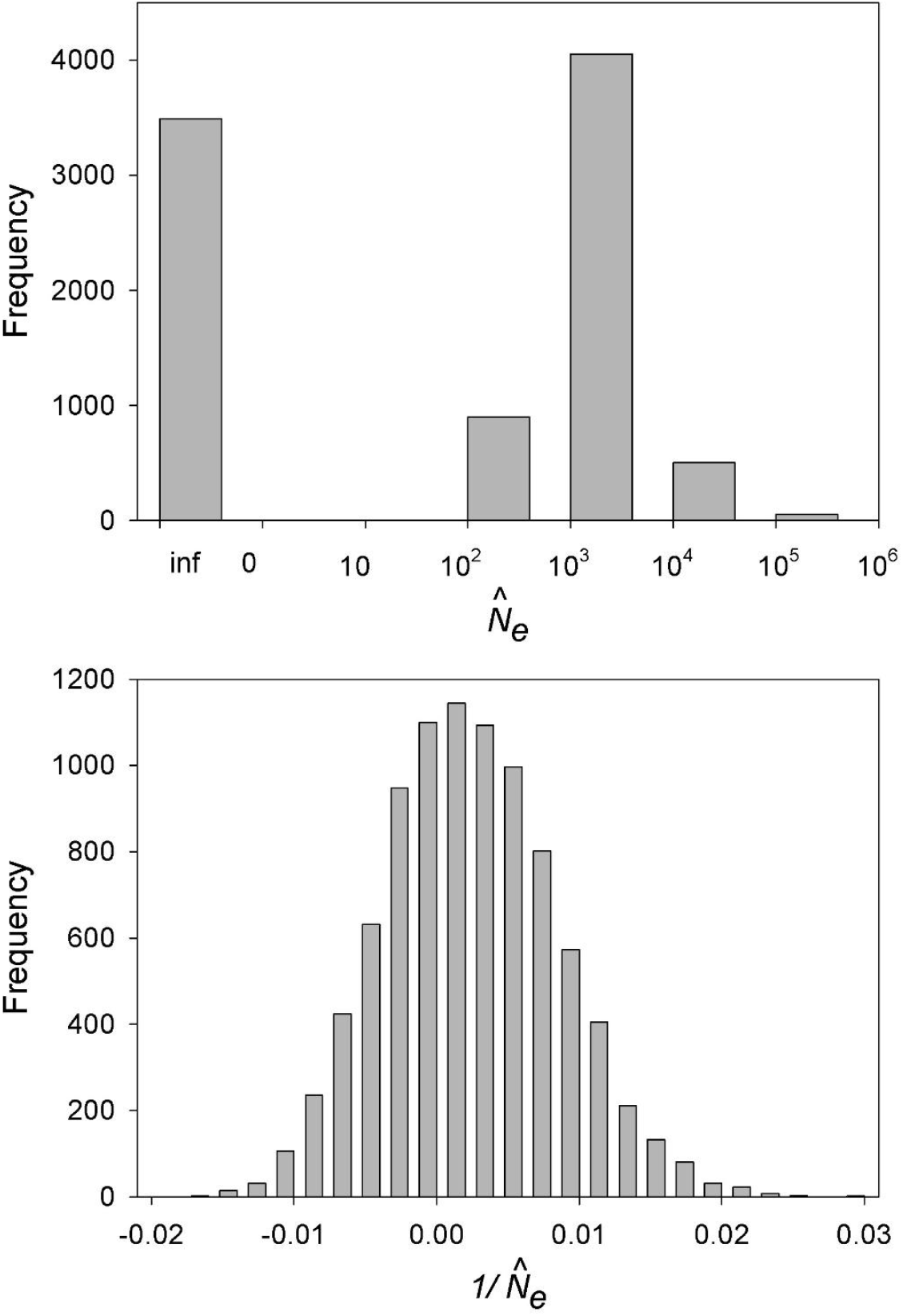
Distribution of 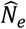 (top) and 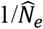 (bottom) in simulated data using the temporal method. The data are from Figure 6 of Waples (2025) for a scenario in which true *N*_*e*_ was 500 and temporal samples of 50 individuals each were taken one generation apart and scored for 100 diallelic loci. In the top panel, negative 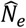 values (farthest left bar) are interpreted as infinity (inf; no evidence for genetic drift). Note the log scale on the X axis of the top panel.

## 2 METHODS

### 2.1 Assumptions

Methods to estimate contemporary *N*_*e*_ generally make the following assumptions, which also are adopted here: generations are discrete; mutation is unimportant over the time frames considered; samples are drawn from a single, isolated population closed to immigration and emigration; and the genetic markers assayed are selectively neutral, such that observed variation reflects the action of random genetic drift. We revisit these assumptions in Discussion.

### 2.2 Expected Values and Variances for Effective Size Estimators

Consider first the standard temporal method, which compares allele frequencies within a single population at two times separated by *t* generations. A widely-used index of change in allele frequency (*P*) is temporal *F* (hereafter just *F*). For a given genetic marker, parametric *F* is the squared difference in population allele frequency between times 0 and *t*, standardized to minimize effects of initial allele frequency:

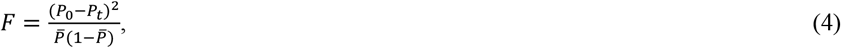

where 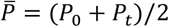 is the mean frequency over time. In general, *P*_0_ and *P*_*t*_ are unknown and must be estimated, from two samples of *S*_*0*_ and *S*_*t*_ individuals. Several authors (Nei and Tajima 1981; Pollak 1983; Jorde and Ryman 2007) have proposed estimators of *F*; differences among these estimators are relatively small and they are collectively referred to here as 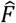. The core premise of the temporal method is that the expected value of 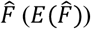 is a simple function of *N*_*e*_ and *t*. Two sampling plans must be considered because they affect the magnitude of allele-frequency change between the samples (Nei and Tajima 1989; Waples 1989):

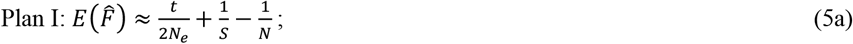

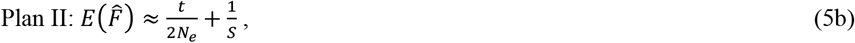

where *N* is census size. The contribution to 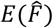 from sampling error is approximately 1/(2*S*_0_) at time 0 and 1/(2*S*_t_) at time *t*. For simplicity, we assume that *S*_*0*_=*S*_*t*_=*S*, so the total magnitude of sampling error is 1/*S*. If *S*_*0*_≠*S*_*t*_, the harmonic mean of the two sample sizes should be used in place of *S*.

In Plan I, individuals for genetic analysis are sampled after reproduction or non-lethally before reproduction; in Plan II, individuals are sampled before reproduction and not allowed to reproduce. The key difference is that under Plan I, some individuals in the initial sample can contribute genes to future generations, which creates positive correlations that have to be accounted for by the 1/*N* term in Equation 5a. Here, *N* is the total number of individuals subject to (presumably random) sampling at time 0; *N* at the time of the second sample is not relevant.

Temporal *F* is closely related to Wright’s (1965) *F*_*ST*_ and related measures of genetic differentiation among populations. Considerable interest has focused on variability of *F*_*ST*_ among loci, as markers with unusually high *F*_*ST*_ might reflect effects of natural selection. Lewontin and Krakauer (1973) proposed that the random drift variance of *F*_*ST*_ among loci is

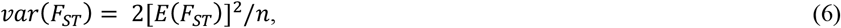

where *n* is the number of independent alleles used to compute each *F*_*ST*_, and that a significantly higher variance than given by Equation 6 could be taken as evidence for the action of selection. Other authors (Robertson 1975; Ewens and Feldman 1976) soon pointed out that various factors besides natural selection can inflate *var*(*F*_*ST*_), and accounting for these factors greatly complicates identification of true outlier loci (Beaumont and Nichols 1996; Lotterhos and Whitlock 2014). The situation is different with temporal *F*, which measures allele-frequency differences within rather than among populations and is not affected by these complicating factors (Gaines and Whittam 1980).

The analogue to Equation 6 that is applicable to the temporal method is

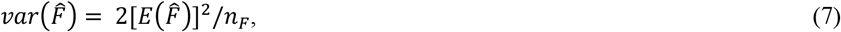

where *n*_*F*_ is the number of independent alleles used to compute 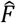. Most genetic datasets now use diallelic SNPs, which have one independent allele per locus, so for our purposes *n*_*F*_ = *L* = number of loci used to compute 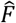. Loci on the same chromosome do not assort independently, and as the number of loci per chromosome becomes large, it becomes more and more likely (and eventually inevitable) that any new locus will be close to, and provide largely redundant information for, one or more existing loci. This lack of independence, or pseudoreplication, reduces the total information content of large datasets. This lack of independence can be quantified by *n’*_*F*_, which is the effective degrees of freedom associated with 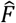. The method described by Waples et al. (2022) was used to estimate *n’*_*F*_ as a function of *L, S, N*_*e*_, and genome size.

In equations 5a and 5b, 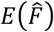 has two components: a drift signal (*t*/(2*N*_*e*_) for Plan II and *t*/(2*N*_*e*_) – 1/*N* for Plan I) and noise from sampling error (1/*S* for both plans). The standard temporal estimator of effective size is obtained by rearranging these equations as follows (Waples 1989):

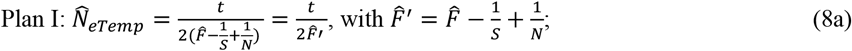

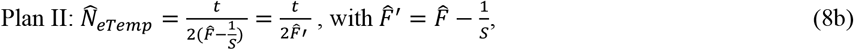

where 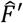 is the raw 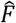 adjusted for sampling error and (for Plan I) the 1/*N* term that accounts for temporal correlations.

Given the above, it is straightforward to derive the theoretical expectation for 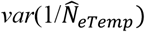:

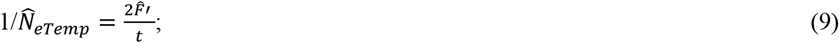

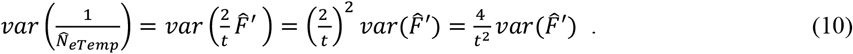

Now adding or subtracting constants does not affect a variance, so 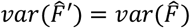, and using the result from Equation 7 for 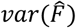,

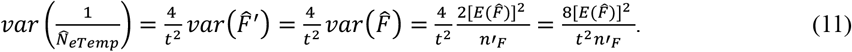

Equation 11 applies to both sampling plans, with the understanding that 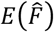 differs between the plans as specified in Equations 5a,b. This means that, all else being equal, 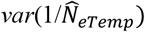 will be smaller for Plan I sampling because 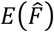 is smaller.

The moment-based LD method, first described by Hill (1981) and reviewed by Waples (2024), has many parallels to the temporal method. The drift signal is captured by *r*^2^, which is the squared correlation of alleles at different loci. If the recombination fraction (*c*) between a pair of loci on the same chromosome is known, the LD method can provide insights into changes of effective size in the past (Tenesa et al. 2007; Hollenbeck et al. 2016; Santiago et al. 2020).

For estimating contemporary *N*_*e*_, focus is on pairs of loci on different chromosomes, which assort independently (*c*=0.5), in which case the estimate applies primarily to the parents of the individuals sampled (Waples 2005). Parametric LD in theory applies to *E*(*r*^2^) in an infinite number of progeny (Weir and Hill 1980). Finite sampling leads to the LD estimator 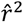, and the Burrows method (Weir 1979; Weir and Hill 1980) is the most widely-used approach for calculating 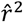. The simplest way to calculate the Burrows coefficient is to use the Pearson product–moment correlation of the genotypic vectors for two loci (Gao et al., 2008).

For unlinked loci, the expectation of 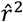 is approximately

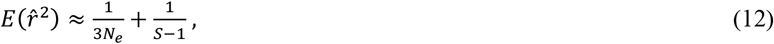

with the first term accounting for genetic drift and the second term for sampling error. Equation 12 is applicable to unphased genotypic data, which is the most common type of data generated by NGS methods for non-model species. Hill (1981) and Weir and Hill (1980) indicated that the contribution to 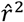 from sampling a finite number of individuals was approximately 1/*S*, but this underestimates effects of sampling error and leads to downward bias in 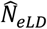 (England et al. 2006; Waples 2006). The term 1/(*S*-1) closely approximates the bias adjustment used in LDNe and implemented in NeEstimator (Do et al. 2014) (unpublished data). Rearranging Equation 12 leads to the LD estimator of contemporary *N*_*e*_

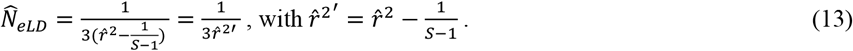

Inverting Equation 13 leads to

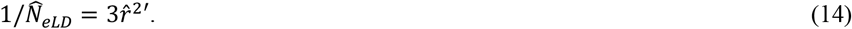

The analogue to Equation 7 that is applicable to the contemporary LD method is

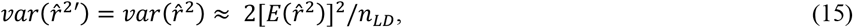

where *n*_*LD*_ is the number of pairs of loci used to compute 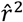. As with the temporal method, physical linkage creates lack of independence for the LD method, but a larger issue is lack of independence caused by overlapping pairs of the same loci. Results from Waples et al. (2022) were used to estimate 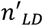, which is the effective number of pairs of loci. Making this substitution, and taking the variance of both sides of Equation 14,

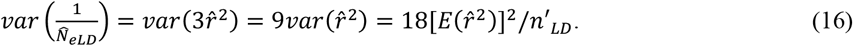

### 2.3 Combining Estimates

Combining two or more estimates can be appropriate when they are estimating the same parameter. Considerations are different for combining estimates within and between methods. In either case, an optimal weighting scheme (in terms of minimizing variance of the combined estimate) accounts for the covariance structure of the estimators (Keller and Olkin 2004). In practice, the covariances will rarely be known. Fortunately, in some important scenarios the estimators can be treated as independent, in which case optimal weights are proportional to reciprocals of variances. In the simulated data, performance of the inverse-variance weighting approach was evaluated under scenarios where the estimators were and were not independent.

#### 2.3.1 Combining estimates within a single method

If multiple samples of individuals provide information about *N*_*e*_ in a single generation for the same population, the simplest approach is to combine the individuals into a single sample. This will reduce the variance of 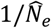 by reducing 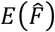 or 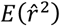 in Equations 5a,b or 12, respectively.

If multiple samples of individuals provide information about *N*_*e*_ in different generations for the same population, using perhaps different sample sizes of individuals and loci, then combining the estimates to increase precision can be appropriate under two scenarios:

- *N*_*e*_ can be considered to be constant over time;
- *N*_*e*_ might vary, but it is of interest to compute a central tendency for *N*_*e*_ over the period encompassed by the samples.

The general approach is the same for these two scenarios (and for both the temporal and LD methods) and is described below.

Assume one has *j*>1 separate estimates of 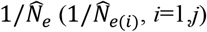, derived from Equations like 9 or 14. Associated with each estimate is a theoretical variance, given by Equations 11 or 16. Denote these variances as *V*_1_, *V*_2_ … *V*_*j*_. The combined estimate 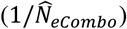 is a weighted mean of the individual estimates, with weights being proportional to reciprocals of variances:

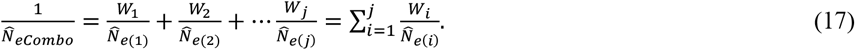

In Equation 17, the weights are standardized such that 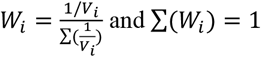.

By extension of Equation (2), the variance of the combined estimate is the sum of the variances plus 2 times all the pairwise covariances:

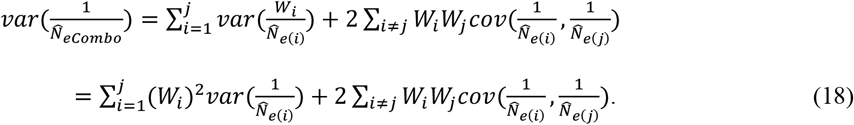

If the different estimates are independent, then the covariance terms drop out and the variance of the combined estimate simplifies to

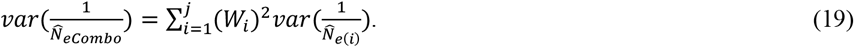

The term *Σ*(*W*_*i*_)^2^ is minimized when all weights are equal.

For the temporal method the raw weights are of the form

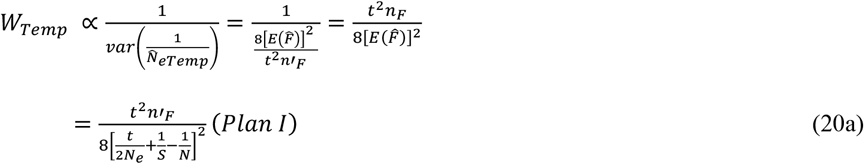

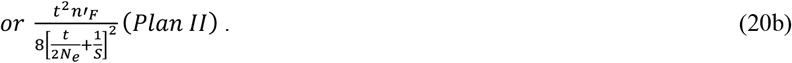

After dividing by the sum of all the raw weights, the standardized weights sum to 1 and can be used in Equation 19. For the LD method, the raw weights are of the form

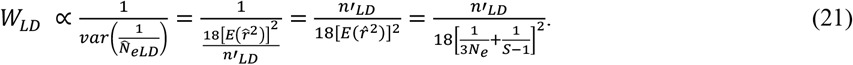

Inspection of Equations 20a,b and 21 indicates that optimal weights for different estimates are functions of true *N*_*e*_, which is unknown. This suggests an iterative process to calculate the weights, as described below.

It is of interest to ask how 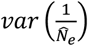 for a combo estimate compares to the variance for a single estimate. For the LD method, if there are *j* estimates that all apply to the same constant or harmonic mean *N*_*e*_, then (assuming constant *S* and 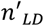) all estimates have the same 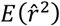 and the same theoretical variance and hence the same weight (*W* = 1/*j*). It follows that

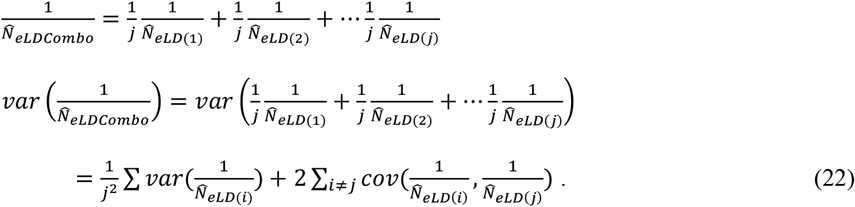

Genetic drift is a Markov process, so it is reasonable to assume independence of LD estimates across time. Ignoring the covariance terms, and substituting for 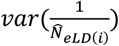 from Equation 16,

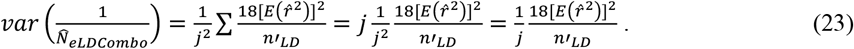

Thus, assuming independence, combining information from *j* equally-weighted LD estimates reduces the overall variance of 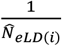 by the factor 1/*j*.

For the temporal method, consider the general case where samples are taken not only in generations 0 and *t*, but also in intermediate generation *t*/2. From Equation 11, the variance of the estimate using the two samples maximally-separated in time is

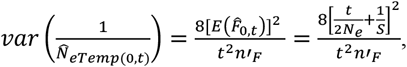

assuming Plan II sampling. The intermediate sample allows one to consider another estimate that combines information from the two shorter estimates:

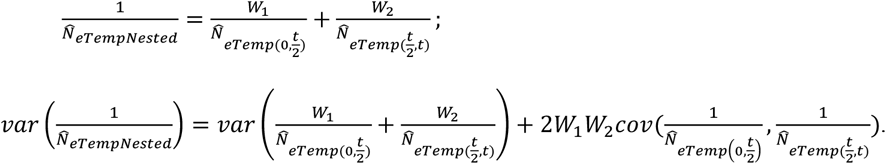

Assuming fixed sample sizes of individuals and loci, *W*_1_ = *W*_2_ = 0.5, and 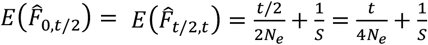. Therefore, letting *C* stand for the covariance term,

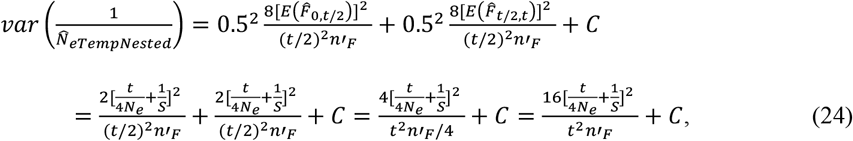

which can be compared with the variance for an estimate based on 2 samples taken *t* generation apart (Equation 11).

#### 2.3.2 Combining estimates across methods

Combining estimates from different methods might be appropriate under either of the scenarios described above for combining estimates within methods. However, probably the greatest benefits from using different methods arise when they estimate *N*_*e*_ in the same time periods, in which case no assumptions are necessary regarding whether *N*_*e*_ varies over time and, if so, to what degree. Care must be taken to ensure that the different methods are estimating *N*_*e*_ in the same generation(s), following rules outlined in Waples (2005). A combined estimate using information from both the temporal and LD methods is of the form

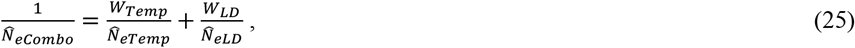

where 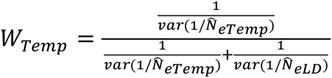 and 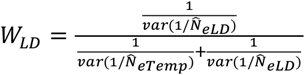.

Comparing the variances gives insight into the relative precision of the two methods:

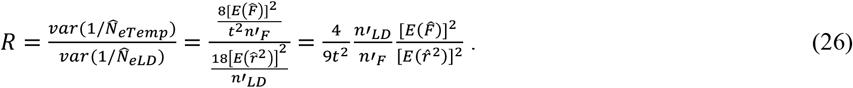

Assuming Plan II sampling for the temporal method, this becomes

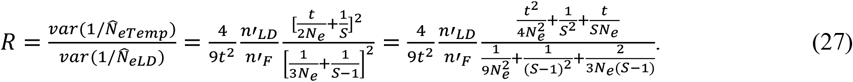

When 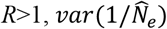 is larger for the temporal method (so the LD method is more precise), and the opposite is true for *R*<1.

More generally, assume that *X* and *Y* both are unbiased estimators of the same parameter *θ*, and further assume that the variance of *X* (*V*_*1*_) is equal to or less than the variance of *Y*. It follows that *var*(*Y*) = *KV*_*1*_, where *K*≥1. Then the standardized weight for *X* is *W*_*1*_ = (1/*V*_*1*_)/(1/*V*_*1*_ + 1/(*KV*_*1*_)) = (1/*V*_*1*_)/((*K*+1)/*KV*_*1*_) = *K*/(*K*+1) and *W*_*2*_ = 1- *W*_*1*_ = 1/(*K*+1). Then, if *X* and *Y* are uncorrelated,

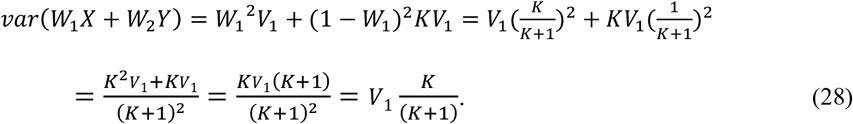

In the special case where the two variances are equal, *K*=1 and *K*/(*K*+1) = 0.5, so the variance of the combined estimate is reduced by 50% compared to either individual variance, as in the example in the Introduction. If *V*_*1*_<*V*_*2*_, the reduction is <50% compared to the smaller variance (Figure S1), and the variance reduction is further restricted if the different estimators are positively correlated.

### 2.4 Computer Simulations

Computer simulations were used to empirically evaluate theoretical results. R code to conduct the simulations will be posted on Zenodo on acceptance. In all simulations, Wright-Fisher populations with constant *N*=*N*_*e*_ were modeled. Populations were initiated with a flat distribution of allele frequencies in the range 0.1-0.5 at a variable number of diallelic (SNP) loci. Within each of 100-1000 replicates, within-method simulations for the temporal method modeled 10 generations of genetic drift. At each generation from *t*=0 to *t*=10, 10 replicate samples of *S* individuals were taken for genetic analysis, according to either Plan I or Plan II protocols. All of these replicate samples were based on the same population pedigree, so they avoided adding spurious demographic variance as described by Waples and Faulkner (2009). Within each replicate, the full matrix of pairwise comparisons of samples was used to generate the full matrix of 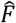 values (for comparisons of generations 0 and 1, 0 and 2, 1 and 2 … 0 and 10). For each type of comparison, means and variances of 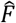 (and the resulting 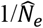 values) were computed across the 10 replicate samples, and these means and variances were then averaged across replicates. Various types of within-method combined estimates were computed as described in the text.

The procedure for within-method LD simulations was similar, except that a 7-generation burnin period was added at the start of each replicate to allow an approximately stable amount of LD to accumulate. The 10 generations of samples were not compared with each other, as in the temporal method; instead, data for each generation were analyzed separately. For each generation, means and variances were computed across the 10 replicate samples, as described above.

Combined estimates across methods are only reasonable when the two methods are estimating the same parameter. Accordingly, these simulations focused on a scenario where two temporal samples of progeny from generations *t* and *t* +1, and a single LD sample of progeny from generation *t* +1, are all used to estimate *N*_*e*_ in generation *t* +1. After initialization and a 7-generation burnin, an initial temporal sample was taken, followed by a single generation of genetic drift and a second temporal sample, which was also used for the LD estimate. As described above, the samples were replicated and means and variances computed across sample replicates and then averaged across replicate simulations.

Correlations between estimates were evaluated three ways. For the temporal and LD methods, correlations were calculated between simulated data series for both within-method and between-method evaluations. Second, simulation code made available in Supporting Information for Waples (2021) was used to quantify correlations between LD and sibship estimates. These simulations assumed that all sibship reconstructions were done without error, so results for the sibship method are optimistic except perhaps for large datasets using ≥1000 SNPs. Finally, the R function MASS::mvrnorm was used to simulate random normal data with specified means, variances, and correlations.

In addition to empirical variances, which measure precision, two more performance metrics were used with the simulated data. The root-mean-squared error (RMSE; square root of the mean squared deviation from the true value) combines information about both precision and bias. Mathematically, RMSE can be calculated as 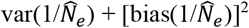. Confidence interval coverage records the percentage of replicates for which the 95% confidence interval around 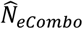 contained the true value of *N*_*e*_. Confidence intervals assumed that 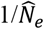 was normally distributed, so the 95% CIs were 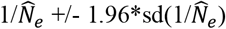.

## 3 RESULTS

I begin by summarizing some of the main analytical results from Methods, followed by presentation of simulation results.

### 3.1 Var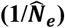

For the temporal method, 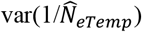 is proportional to squared 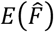 (Equation 11), which varies according to the sampling plan as shown in Equations 5a,b. All else being equal, 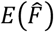 is smaller for Plan I sampling by the magnitude 1/*N*, so Plan I sampling leads to higher precision. 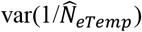 is reduced by the number of generations between samples and by the effective number of alleles used in the calculation of 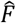 (Equation 11). In simulations, Equation 11 accurately predicted 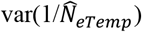 for samples spaced 1-10 generations apart (Figure S2, top panel).

For the LD method, 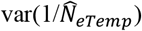 is proportional to squared 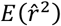 and it is inversely proportional to the number of effectively independent pairs of loci used in the calculation of 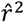 (Equation 16). In the simulations, Equation 16 tended to slightly underestimate (by a few percent) 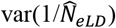 for single samples of progeny (Figure S2, bottom panel).

### 3.2 Combining Estimates Within Methods

Before combining multiple estimates of 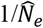, it is necessary to calculate appropriate weights, which should be standardized so they sum to 1. We use weights that are inversely proportional to 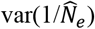, so temporal weights (Equations 20a,b) are inversely proportional to Equation 11 and LD weights (Equation 21) are inversely proportional to Equation 16.

Standardized weights are of the form

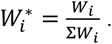

Per Equation 2, variances of combined estimates are the sum of the individual weighted variances, plus covariance terms for each pair of estimates being averaged. Single-sample estimates are simpler to analyze in this regard because they are arguably independent across generations, in which case the covariance terms vary randomly around 0 and can be ignored. If so, the combined LD estimate that leverages information from separate LD estimates in *j* generations has expected variance that is (1/*j*) times the variance of an estimates based on a single-generation of data (Equation 23). Simulation results (Figure S3, top panel) were in excellent agreement with this prediction, indicating that it is reasonable to ignore the covariance terms in 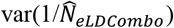. RMSE of the combined LD estimates also declined systematically as information from more samples was utilized (Figure S3, bottom panel).

Combining multiple temporal estimates is trickier because the estimates typically are positively correlated. Two general scenarios can be considered. In the first (nested) scenario, a single pair of samples from generations 0 and *t* is supplemented by an additional sample at generation *t*/2. 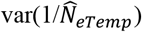 for the two extreme samples is given by Equation 11. The other two pairs of samples each span half the total time (*t*/2 generations) and have the same weights and the same 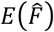, and the combined variance is given by Equation 24. Even ignoring the covariance term, 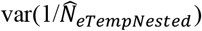 is generally larger than the variance from the two samples spanning the full *t* generations (Equation 11), unless *S* is relatively large compared to *N*_*e*_ and/or *t* is very large (Table S1). In this nested scenario, however, the covariance term will be positive because the two estimates based on the shorter time frame share the same sample at generation *t*/2. This covariance will become relatively less important as *t* increases. In the simulations, 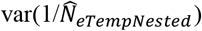 was markedly higher than the single estimate for *t*=2 and only modestly higher for *t*=10 (Figure S4, top). Thus, splitting a long temporal comparison into two shorter ones does not improve precision.

In the second scenario, all 3 estimates (two spanning *t*/2 generations and one spanning all *t* generations) are combined into a single, comprehensive estimate, using weights calculated as in Equations 20a,b. The estimates spanning short and long time periods are highly positively correlated, as they not only share samples but also duplicate coverage of overlapping generations of genetic drift. These positive correlations increase the variance of the comprehensive estimates and limit the benefits of adopting this scenario. Nevertheless, simulations showed that precision did increase somewhat by using all of the data (compared to the single estimate, 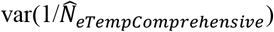 was reduced by 13% for *t*=2, 23% for *t*=5, and 24% for *t*=10; Figure S4, bottom).

### 3.3 Combining Estimates Across Methods

#### 3.3.1 Temporal and LD methods

The general formula for combining estimates across temporal and LD methods is (Equation 22)

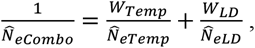

where the raw weights are calculated as in Equations 20 and 21 and standardized weights are of the form

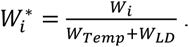

It is useful to express the theoretical variances of the two methods as a ratio (*R*), which provides insights into relative precision (Equation 26):

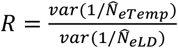

*R*>1 indicates higher variance (less precision) for the temporal method, and *R*<1 indicates the reverse. If temporal estimates span a single generation (*t*=1), the LD method has greater precision unless at least 10^4^ SNPs are used (Figure 2, top). Datasets in the range of 100-1000 SNPs are in the “sweet spot” for the LD method, where the benefits of being able to use information from many pairwise combinations of loci are maximized. As more and more SNPs are used, lack of independence limits further increases in precision more for the LD method than it does for the temporal method. All else being equal, relative precision of the LD method is higher when true *N*_*e*_ is larger. When temporal estimates span multiple generations (*t>*1), relative precision of the temporal method increases proportionally, resulting more frequently in *R*<1 (Figure 2, bottom). In this scenario, however, a single LD estimate applies to only one of the (potentially many) generations over which *N*_*e*_ is estimated by the temporal method, so that must be kept in mind in interpreting what 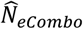 represents.

**Figure 2.**
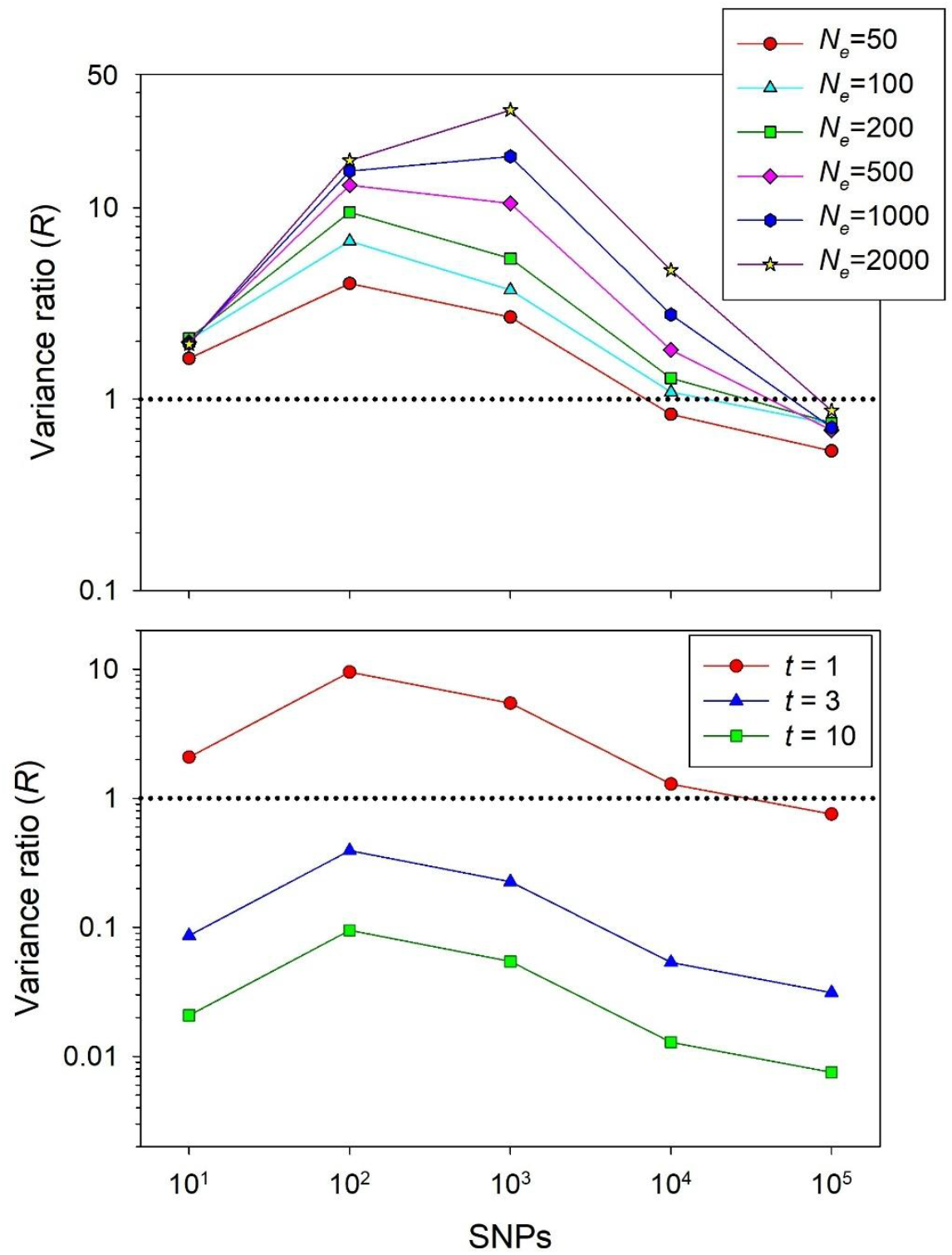
Relative precision of the temporal and LD methods for estimating effective size. The Y axis shows theoretical ratio (*R*) of 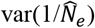 for the temporal method to 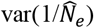 for the LD method, as a function of true *N*_*e*_, the number of SNPs used to estimate effective size, and elapsed time between temporal samples. Temporal estimates used 2 samples of 50 individuals; LD estimates used one sample of 50 progeny. Top: Effects of *N*_*e*_, with all temporal samples taken 1 generation apart. Bottom: results for temporal samples taken *t* = 1, 3, or 10 generations apart, with *N*_*e*_ fixed at 200. *R* was computed using Equation 23, assuming Plan II sampling for the temporal method. The horizontal dotted line indicates equal variances for the two methods. These results account for pseudoreplication (lack of independence) caused by physical linkage and (for the LD method) overlapping pairs of the same loci, as quantified by Waples et al. (2022). These results assume 16 chromosomes of 50Mb (mean chromosome numbers for plants, invertebrates, and vertebrates are 13, 11, and 25, respectively; Li et al. 2011). Note the log scale on both axes.

The scenario with *t*=1 is of particular interest because in that case both the temporal and LD methods estimate effective size in a single generation, which facilitates estimation of the key ratio *N*_*e*_/*N*. Fortunately, the two estimators are effectively uncorrelated, even for relatively large values of *L* and *S* (Table 1). This means that the covariance term can be ignored in estimating 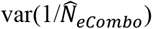. For the scenarios considered in Table 1, the LD method was always more precise (smaller 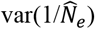) than the temporal method, but properly combining the two estimates still provided benefits (reduced 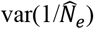 and RMSE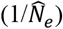 for 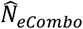; Tables 1C and 1D).

**Table 1.** Performance metrics for combined estimates based on the LD and temporal estimators. Results are for simulations using sample sizes of individuals (*S*) ranging from 25 to 100 and numbers of unlinked, diallelic (SNP) loci ranging from 100 to 5000. True *N*_*e*_ was 200. A: Correlations between 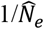 for the two methods. B: Confidence interval coverage—percentage of replicates for which the 95% confidence interval around 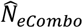 contained the true value. C: ratio of 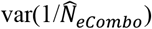 to 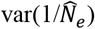 for LD method. D: ratio of the root-mean-squared error (RMSE) of 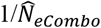 to 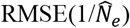 for LD method. For the scenarios modeled, 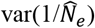 and 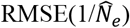 were both smaller for the LD method than for the temporal method.

| A | Number of loci |  |  |  |
| --- | --- | --- | --- | --- |
|  | S | 100 | 1000 | 5000 |
|  | 25 | -0.005 | -0.007 | 0.004 |
|  | 50 | 0.000 | -0.006 | -0.002 |
|  | 100 | -0.010 | 0.004 | 0.002 |

| B | Number of loci |  |  |  |
| --- | --- | --- | --- | --- |
|  | S | 100 | 1000 | 5000 |
|  | 25 | 94.8 | 94.8 | 73.1 |
|  | 50 | 96.3 | 94.0 | 74.5 |
|  | 100 | 99.1 | 99.2 | 95.1 |

| C | Number of loci |  |  |  |
| --- | --- | --- | --- | --- |
|  | S | 100 | 1000 | 5000 |
|  | 25 | 0.94 | 0.87 | 0.62 |
|  | 50 | 0.94 | 0.89 | 0.65 |
|  | 100 | 0.95 | 0.96 | 0.87 |

| D | Number of loci |  |  |  |
| --- | --- | --- | --- | --- |
|  | S | 100 | 1000 | 5000 |
|  | 25 | 0.97 | 0.92 | 0.76 |
|  | 50 | 0.97 | 0.94 | 0.79 |
|  | 100 | 0.97 | 0.98 | 0.92 |

The benefit of combining estimates is maximized when they are uncorrelated and have equal variances (Equation 28). This is illustrated in Figure 3, which shows how 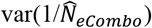 varies in simulated data as a function of the weight assigned to the LD method (*W*_*LD*_). In the top panel (using 5000 SNPs), the optimal weights are *W*_*LD*_ = 0.63 and *W*_*Temp*_ = 0.37 and the optimal combined estimate has a variance that is 37% lower than that of the LD estimates alone (and 63% lower than the variance of the temporal method alone). Conversely, if the variances are very different, the benefits of combining estimates can be marginal at best (maximum improvement <10% for the scenario in bottom panel of Fig 4, where 500 SNPs are used, which is in the “sweet spot” for relative precision of the LD method).

**Figure 3.**
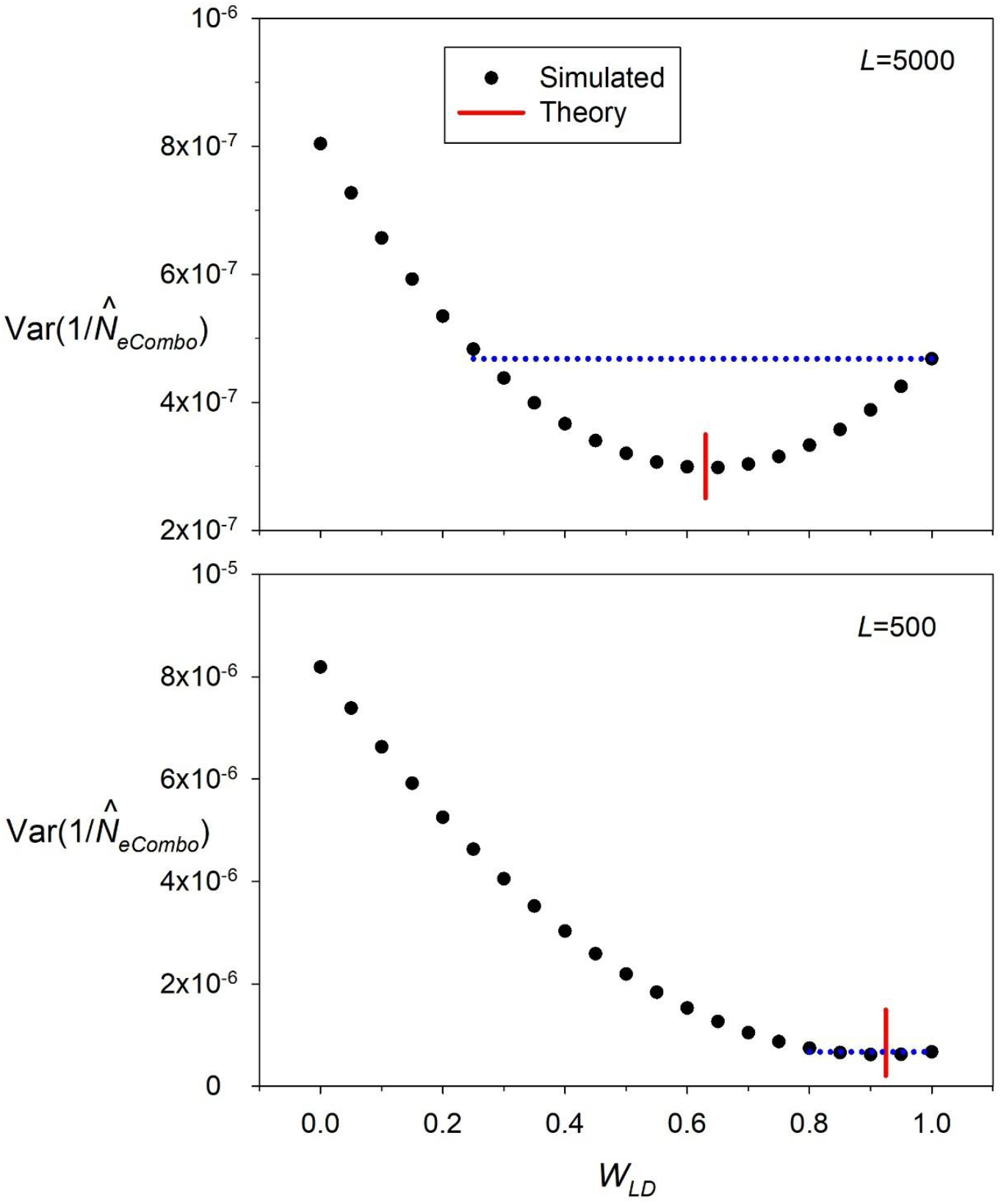
Variance of a combined estimate (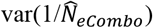; Y axis), as a function of the relative weight for the LD method (*W*_*LD*_; X axis). Combined estimates were computed as 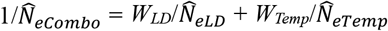, where *W*_*Temp*_ = 1-*W*_*LD*_. Filled circles are results from simulations; vertical lines show the theoretical optimum weighting scheme based on Equations 11, 16, and 28. Horizontal dotted lines show the range of values of *W*_*LD*_ that produce 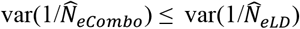. Simulation parameters were *N*_*e*_ = 200, *S* = 50, *t*=1 for the temporal method, and using *L*=5000 (top) or 500 (bottom) unlinked, diallelic loci.

**Figure 4.**
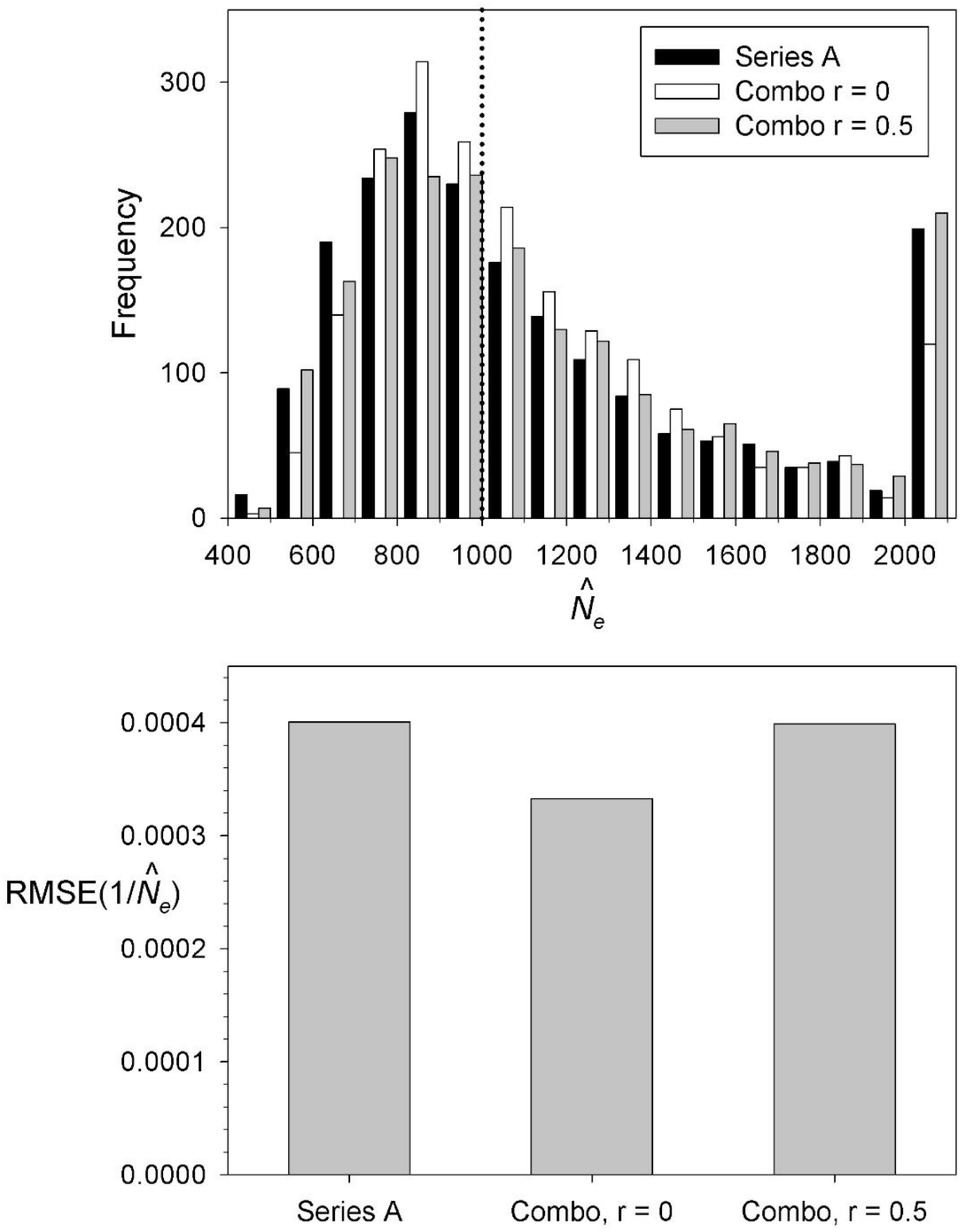
Top: Distribution of 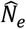 in random simulated data. Two series of 2000 paired, random normal datapoints were simulated. Both series (A and B) had an expected value of 1/*N*_*e*_ = 1/1000, and the variance of B was twice that of A. A third, combo series 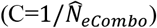 was generated as the weighted average of A and B, with weights of 0.67 and 0.33, respectively, based on reciprocals of variances. The inverse of the simulated data are the 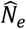 values shown in the graph, with true *N*_*e*_ (1000) indicated by a dotted vertical line. The last group of bars on the right include all 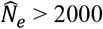. Black bars show the distribution of Series A. White bars show the distribution of 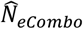 when A and B are modeled as uncorrelated (realized correlation coefficient = −0.02), and gray bars show the distribution of 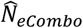 when A and B are modeled with expected correlation 0.5 (realized correlation coefficient = 0.51). Bottom: RMSE of 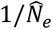 for Series A and for 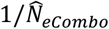 in the uncorrelated and correlated scenarios.

Figure 3 also illustrates the consequences of improperly weighting the individual estimates. In the top scenario, 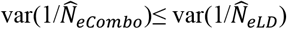 as long as *W*_*LD*_ is anything larger than about 0.25. But in the bottom scenario, with very skewed weights, 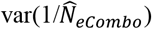 can be greater than 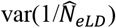 unless the weight assigned to the LD estimate is 0.8 or larger.

If information from two methods is being combined and both methods use information from multiple samples, estimates within methods should be combined first, and an overall estimate of 1/*N*_*e*_ can be computed as a weighted mean of the two combined within-method estimates. Note that because of positive correlations among nested estimates, estimating the variances associated with comprehensive temporal estimates can be challenging.

Effects of positive correlations are illustrated with simulated data in Figure 4. Using a target *N*_*e*_ of 1000, two series of random normal variates were simulated, both having expected value = 1/*N*_*e*_ = 1/1000. The variance of series *B* was twice that of series *A*, so *A* was weighted more heavily in computing the combined estimate 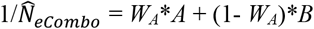. When the two series were simulated independently (white vertical bars in top panel in Figure 4), the distribution of 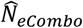 was considerably tighter than for series *A* (black bars) or *B* (not shown), with fewer estimates >2x the true value. But when the two series were simulated jointly with expected positive correlation of 0.5 (gray bars), computing a combined estimate led to little or no increase in precision compared to that provided by Series *A* alone.

#### 3.3.2 The sibship method

The other widely-used single-sample estimator is the sibship method developed by Wang (2009). Analyses directly comparable to those described above for the temporal and LD methods are difficult because no analytical formula for 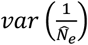 analogous to Equations 11 or 16 exists for the sibship method. Instead, uncertainty in *N*_*e*_ estimation has been handled in two ways. For individual point estimates, confidence intervals can be generated as part of the maximum-likelihood estimation process (Wang and Santure 2009). Second, for selected simulation scenarios, Wang (2009, 2016) has published empirical variances of 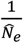 or RMSE values. Wang’s comparisons have shown that a) precision of the sibship and LD methods is much higher than for either the coancestry or heterozygtote excess methods, and b) precision generally was a bit higher for the sibship than the LD method, but the difference narrowed as more loci were used. Most of these comparisons involved relatively small numbers of loci (<100).

Unlike the temporal and LD methods, the sibship method has a hard upper limit to performance, because once enough loci are used to correctly identify all siblings within a sample, adding more loci cannot further increase accuracy or precision. Waples (2021) showed that, because of this limitation, in genomics-scale datasets (≥1000 SNPs) precision of the LD method will generally exceed that of the sibship method, even if all siblings are correctly identified. Waples (2021) also examined the correlations between 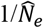 for the sibship and LD methods and found that correlation coefficients were generally low for small datasets but could be very high for genomics-scale datasets. Typical results are shown in Table 2: correlations were generally <0.1 for 100 or fewer loci but could be as high as 0.9 when more than a thousand SNPs were used. For the same number of loci, correlations were generally lower for smaller sample sizes of individuals. For example, with true *N*_*e*_=200 and using at least 1000 SNPs, Pearson’s *r* was 0.79 or higher for *S*=50 or 100 but ≤0.21 for *S*=25.

**Table 2.** Correlations between 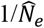 for the sibship and LD methods in simulated data. Simulations used the software that is available in Supporting Information for Waples (2021), which assumes that all sibling assignments are made without error. True *N*_*e*_ was 200 (top) and 2000 (bottom). *S* is the number of individuals sampled.

| $N_e=200$ | | Number of loci | | | |
| --- | --- | --- | --- | --- | --- |
| $S$ | 10 | 100 | 1000 | 2500 | |
| 25 | 0.024 | 0.085 | 0.171 | 0.211 |  |
| 50 | 0.034 | 0.089 | 0.844 | 0.892 |  |
| 100 | 0.029 | 0.306 | 0.791 | 0.839 |  |
| $N_e=2000$ | | Number of loci | | | |
| $S$ | 10 | 100 | 1000 | 2500 | |
| 25 | 0.011 | 0.117 | 0.335 | 0.183 |  |
| 50 | -0.011 | 0.060 | 0.302 | 0.285 |  |
| 100 | 0.001 | 0.032 | 0.730 | 0.904 |  |

### 3.4 Confidence Intervals

Two major factors can degrade performance of confidence intervals: 1) incorrectly calculating the variance of the estimator; 2) using a biased estimator. Superior performance occurs when all estimators are unbiased and variances are correctly estimated. Both of these conditions were met for the combined estimates shown in Figure 4 when Series A and Series B were simulated independently (white vertical bars). 95.0% of the nominally-95% CIs included the true *N*_*e*_, 2.6% of the CIs were too low, and 2.4% were too high (first row of data in Table S2). Results were similar when Series A and Series B were simulated with expected correlation 0.5 (gray vertical bars) and the expected variance of the combined estimate was computed using Equation 3 to account for the resulting covariance term (second row of data in Table S2). But when correlated data were simulated but the resulting covariance was ignored in computing the estimated variance, the CIs around 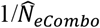 were too narrow and included the true value less than 90% of the time (third row of data in Table S2).

The effect of bias on performance of confidence intervals can be seen for within-method analyses using the LD method. For parameter values used in the simulations, 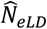 has an upward bias of about 5% (Figure S2), which has been documented before (Waples and Do 2010) and is the result of a slight over-correction for downward bias in the original version (Hill 1981) of the LD method. When only a single estimate is considered, based on 50 individuals scored for 1000 SNPs, this modest bias has little effect on CI performance (93.8% coverage; first line in Table S3). But when multiple samples are used and the width of the CIs shrinks, coverage declines. When information from 10 LD samples is combined, coverage has dropped to 80.7%, and almost all CIs that don’t include the true *N*_*e*_ are too high (last line of Table S3). This latter result occurs because as the estimates become more precise and the width of CIs shrinks, they converge on a value larger than the true *N*_*e*_. An estimator with a downward bias would have the opposite effect: CIs that did not contain the true *N*_*e*_ would tend to be too low.

Table S3 shows that in the presence of bias, CI performance degrades as more samples are combined to increase precision. A similar phenomenon occurs when one increases the number of loci. For the analyses reported in Table 1, an LD estimate based on a single sample was combined with a temporal estimate based on two samples spanning a single generation. With only 100 loci, coverage of the CIs for the combined estimates was actually >95% (Table 1D), despite the bias contributed by the LD method. But when 5000 loci were used, coverage dropped below the expected 95%, and substantially so for the smaller sample sizes.

### 3.5 Details Regarding Estimation

Combining multiple estimates to arrive at a single estimate of effective size involves the following general steps:

1. Generate 2 or more individual estimates of 1/*N*_*e*_;
2. Estimate the variance associated with each estimate;
3. Assign weights based on reciprocals of the variances from Step 2;
4. Compute the weighted mean of the two estimators to obtain 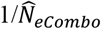;
5. Generate CIs for 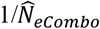;
6. Invert the values in Step 5 to generate the point estimate 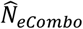 and its CIs.

In Step 2, variances for the LD and temporal methods can be estimated using Equations 11 and 16, respectively. Alternatively, if a series of estimates is available for a single method, the empirical variance of 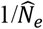 could be used. Step 3 uses Equations 20-21 to compute weights, and Step 4 uses the generalized Equation 17 to calculate 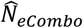. Step 5 takes advantage of the fact that 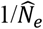 will be approximately normally distributed as long as the number of datapoints used in its calculation is at least 50. In contemporary datasets, 50 loci or independent alleles is generally easy to achieve for the temporal method and even easier for the LD method, which uses pairs of loci.

Optimal weights for 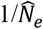 for the temporal and LD methods depend on the respective variances, which are functions of true *N*_*e*_ (cf Equations 11 and 16). As true *N*_*e*_ is unknown, an iterative process is used:

1. Compute an initial 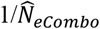 using equal weights;
2. Compute initial weights assuming true 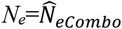;
3. Repeat Steps 1 and 2 until convergence, using the updated weights in Step 1 and the updated 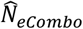 in Step 2.

Evaluations indicate that 3 iterations is sufficient for convergence of most scenarios, but some can require 5 or more.

### 3.5 Software

*ComboNe*, coded in R, is introduced to carry out analyses described here. The R code, together with a sample input file and a ReadMe text file, is available in Supporting Information.

### 3.6 A Worked Example

A worked example uses the following input data for the temporal and LD methods (which are the first row of data from the sample input file): *L* = 9100 SNPs; *S* = 40 individuals; 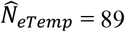 and 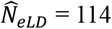. The initial equally-weighted estimate of *N*_*e*_ is 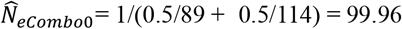. Properly weighting the two estimates requires knowing the effective numbers of independent datapoints (effective degrees of freedom) associated with each. Embedded in *ComboNe* is code from Waples et al. (2022) that estimates *n’*_*LD*_ and *n’*_*F*_ as a function of *N*_*e*_, *S*, genome size, and *L*. For illustration, we assume the focal species has 16 chromosomes, in which case the initial effective degrees of freedom estimates are *n’*_*F*_ = 3118 for the temporal method and *n’*_*LD*_ = 7940 for the LD method. Based on Equation 11, the initial estimate for 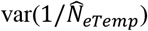 is *V1*_*Temp*_ = (8*(1/(2\**N*_*e*_) + 1/*S*))/*n’*_*F*_. Substituting the above values, *V1*_*Temp*_ = (8*(1/(2*99.96) + 1/40))/3118 = 7.698×10^−5^ and inverting that gives raw *W1*_*Temp*_ = 12991. Similarly, using Equation 16, the initial estimate for 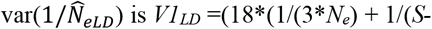 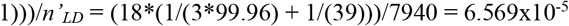, and inverting that gives the initial raw weight of 15223. After standardization so that the weights sum to 1, the standardized weights are *W1\**_*Temp*_ = 0.460 and *W1\**_*LD*_ = 1-0.460 = 0.540. With the first estimated weights, we update the combined estimate of *N*_*e*_ as 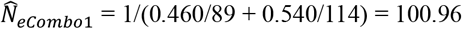. In the second iteration, the estimated variances and effective degrees of freedom are updated with the new 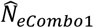, leading to very slightly different weights (*W2\**_*Temp*_ = 0.461 and *W2\**_*LD*_ = 0.539.) The new weights change only the third decimal place of 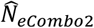, and the third iteration produces little more in the way of change. So the best combined estimate is 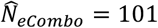, with the two methods providing roughly equal amounts of information.

The final step is to calculate CIs for the combined estimate. After the third iteration, the final estimates of 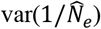 were 2.302×10^−6^ for the temporal method and 1.899×10^−6^ for the LD method. From Equation 19, assuming independence (as is appropriate for temporal/LD combinations), 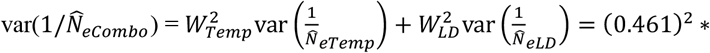 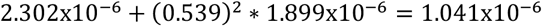, so the standard deviation of 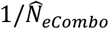 is 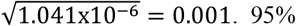 CIs for 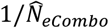 are thus 1/101 +/-1.96*0.001 = 0.0099 +/-0.00196 = [0.00794, 0.01186]. Inverting these gives the upper 95% CI for 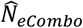 as 126 and the lower CI as 84. With this much data, and combining information from two independent methods, it is possible to make fairly precise statements about effective population size.

## 4 Discussion

Using genetic methods to estimate effective populations size has become something of a cottage industry—and a fairly large one. It is relatively easy to use the same or overlapping datasets to estimate effective size using more than one method, and many authors do so, but it remains relatively rare to report combined estimates that leverage information from more than one method. This is unfortunate, as combining estimates can substantially improve precision, which is often the limiting factor in *N*_*e*_ estimation.

The approach outlined here improves on previous approaches by focusing on the variance of 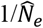 rather than the variance of 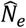, which is problematical because the distribution of 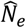 is highly non-normal. 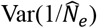 is closely related to the variances of temporal 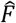 and 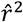, and the distributions of these metrics are well understood and effectively normal if the number of independent datapoints used in their calculation is larger than about 50. As a consequence, it is straightforward to calculate the variance of a combined estimate 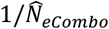 and use that to set realistic confidence intervals for 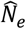 using standard statistical theory. Benefits of doing this include fewer very high or infinite estimates and tighter confidence intervals with higher lower bounds, all of which improve the value of effective size estimates for applied conservation and management.

A precondition for combining estimates is that the different methods must be estimating the same parameter. This can potentially be appropriate either in combining estimates for different time periods within methods (see Section 2.3.1 for details) or for combining estimates for different methods for the same time period(s) (see Section 2.3.2 for details). The latter scenario is particularly well-suited to increasing precision of the estimated *N*_*e*_/*N* ratio.

Benefits of combining data from multiple estimators are maximized under three general conditions:

1. The variances (and hence weights) of individual estimators are equal. In this scenario, 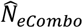 is the unweighted harmonic mean of the individual 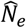 values.
2. The estimators are uncorrelated. Strictly speaking, precision is maximally increased when the estimators are negatively correlated, but that is not likely to occur in real datasets. Therefore, in practical terms, the ideal situation occurs when positive correlations are low or non-existent.
3. Individual estimates have low precision, which occurs when data are limited and/or true *N*_*e*_ is large.

Condition 1 (equal variances) is ideal because the reduction in variance of the combined estimate diminishes as the weights become more uneven (Equation 28; Figure S1). When weights are very unequal, most of the useful information is provided by just one of the methods and combining them provides relatively little benefit (Figure 3, bottom panel). The temporal and LD methods make it easy to properly weight combined estimates because 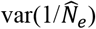 is straightforward to estimate. At present there is not a simple, generic way to calculate appropriate weights for Wang’s sibship method. For some specific parameter combinations, Wang (2009, 2016) and Waples (2021) have provided estimates of 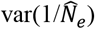 or related quantities. In other scenarios, giving equal weights to sibship and LD estimates might be reasonable as a first approximation.

Condition 2 (uncorrelated estimators) appears to be a valid assumption for the following scenarios: (a) combining temporal and LD estimators for the same time periods; (b) combining LD estimates for different time periods; (c) combining LD and sibship estimates when *N*_*e*_ is not too small and relatively few genetic markers are used. Although I am not aware of a direct evaluation, it seems likely that estimates from the temporal and sibship methods are also essentially uncorrelated. When the number of genetic markers is large (≥1000 SNPs), both the LD and sibship methods become very good at detecting essentially the same signal of genetic drift, with the consequence that positive correlations reduce the benefits of combining estimates. Positive correlations also limit benefits from combining temporal estimates based on overlapping samples and/or time periods, but this is a complex topic that merits further study. One common scenario has already been examined in detail. If a time-series of consecutive samples is taken as part of a systematic genetic monitoring program, precision of temporal estimates of *N*_*e*_ in individual generations can be improved by using information from comparisons than span one additional generation on either side of the focal generation (using the program *M**ax**T**emp* described by Waples et al. 2025).

Condition 3 represents good news for researchers, as it means that the greatest improvements to precision can be achieved when they are needed the most.

All of the above analyses have focused on precision. Most population genetics theory begins with simplifying assumptions to make analyses tractable (models for estimating effective size are no exception), and violations of these core assumptions can lead to bias. Two of the most consequential assumptions for *N*_*e*_-estimation methods are that (1) generations are discrete, and (2) the focal population is isolated. Both of these assumptions are commonly violated in real populations. Table 3 of Waples et al. (2025) provides a compilation of numerous published papers that have evaluated sensitivities to these and other potential sources of bias, and researchers interested in combining estimates from multiple methods should also consider their respective biases.

Performance of confidence intervals (summarized in Tables 1, S2, and S3) illustrates some potential tradeoffs between precision and bias. If any estimator has even modest bias, the bias will have more pronounced effects on CI coverage as precision increases and CI width shrinks. Thus, it is possible for combined estimates of *N*_*e*_ to have increased precision but lower CI coverage. Researchers should be aware of this possibility and consider an experimental design that maximizes the chances of achieving their objectives.

## Data Accessibility and Benefit-Sharing

### Data accessibility statement

No data were used in this paper except those that a) were generated by simulations, or b) have been previously published. Code to conduct simulations described here will be posted on Zenodo on acceptance.

### Benefit-sharing statement

This is a theoretical paper, so this statement is not applicable.

## References

Beaumont, M.A. and Nichols, R.A., 1996. Evaluating loci for use in the genetic analysis of population structure. Proceedings of the Royal Society of London. Series B: Biological Sciences, 263(1377), pp.1619–1626.

Clarke, S.H., Lawrence, E.R., Matte, J.M., Gallagher, B.K., Salisbury, S.J., Michaelides, S.N., Koumrouyan, R., Ruzzante, D.E., Grant, J.W. and Fraser, D.J., 2024. Global assessment of effective population sizes: Consistent taxonomic differences in meeting the 50/500 rule. Molecular Ecology, 33(11), p.e17353.

Cochran, W. G. (1937). Problems arising in the analysis of a series of similar experiments, Supplement to the Journal of the Royal Statistical Society 4 102–118.

Convention on Biological Diversity (CBD). 2022. “Decision Adopted by the Conference of the Parties to the Convention on Biological Diversity CBD/COP/DEC/15/4 Kunming-Montreal Global Biodiversity Framework. CBD/COP/DEC/15/4. https://www.cbd.int/doc/decisions/cop-15/cop-15-dec-04-en.pdf.

Do, C., R.S. Waples, D. Peel, G.M. Macbeth, B.J. Tillet, and J.R. Ovenden. 2014. NeEstimator V2: re-implementation of software for the estimation of contemporary effective population size (Ne) from genetic data. Molecular Ecology Resources 14:209–214 (DOI: 10.1111/1755-0998.12157).

England, P. R., Cornuet, J.-M., Berthier, P., Tallmon, D. A., & Luikart, G. (2006). Estimating effective population size from linkage disequilibrium: Severe bias using small samples. Conservation Genetics, 7, 303–308.

Ewens, W.J. and Feldman, M.W., 1976. The theoretical assessment of selective neutrality. Population genetics and ecology, pp.303–337. S. Karlin and E. Nevo, eds. Academic Press, NY.

Gaines, M.S. and Whittam, T.S., 1980. Genetic changes in fluctuating vole populations: selective vs. nonselective forces. Genetics, 96(3), pp.767–778.

Gao, X., Starmer, J., & Martin, E. R. (2008). A multiple testing correction method for genetic association studies using correlated single nucleotide polymorphisms. Genetic Epidemiology: The Official Publication of the International Genetic Epidemiology Society, 32(4), 361–369.

Hill, W.G., 1981. Estimation of effective population size from data on linkage disequilibrium. Genetics Research, 38(3), pp.209–216.

Hollenbeck, C., Portnoy, D., & Gold, J. (2016). A method for detecting recent changes in contemporary effective population size from linkage disequilibrium at linked and unlinked loci. Heredity, 117, 207–216.

Jorde, P.E. and Ryman, N., 2007. Unbiased estimator for genetic drift and effective population size. Genetics, 177(2), pp.927–935.

Keller, T. and Olkin, I., 2004. Combining correlated unbiased estimators of the mean of a normal distribution. Lecture Notes-Monograph Series, pp.218–227.

Lewontin, R.C. and Krakauer, J., 1973. Distribution of gene frequency as a test of the theory of the selective neutrality of polymorphisms. Genetics, 74(1), pp.175–195.

Lotterhos, K.E. and Whitlock, M.C., 2014. Evaluation of demographic history and neutral parameterization on the performance of FST outlier tests. Molecular ecology, 23(9), pp.2178–2192.

Mastretta-Yanes, A., J. M. Da Silva, C. E. Grueber, et al. 2024. ”Multinational Evaluation of Genetic Diversity Indicators for the Kunming-Montreal Global Biodiversity Framework.” Ecology Letters 27, no. 7: e14461.

Nadachowska-Brzyska, K., M. Konczal, and W. Babik. 2022. “Navigating the Temporal Continuum of Effective Population Size.” Methods in Ecology and Evolution 13: 22–41.

Nei, M. and Tajima, F., 1981. Genetic drift and estimation of effective population size. Genetics, 98:625–640.

Nomura, T., 2008. Estimation of effective number of breeders from molecular coancestry of single cohort sample. Evolutionary applications, 1(3), pp.462–474.

Palstra, F. P., and D. J. Fraser. 2012. “Effective/Census Population Size Ratio Estimation: A Compendium and Appraisal.” Ecology and Evolution 2: 2357–2365.

Pollak E (1983) A new method for estimating the effective population size from allele frequency changes. Genetics, 104, 531–548.

Pudovkin, A.I., Zaykin, D.V. and Hedgecock, D., 1996. On the potential for estimating the effective number of breeders from heterozygote-excess in progeny. Genetics, 144(1), pp.383–387.

Robertson, A., 1975. Gene frequency distributions as a test of selective neutrality. Genetics, 81(4), pp.775–785.

Santiago, E., Novo, I., Pardiñas, A. F., Saura, M., Wang, J., & Caballero, A. (2020). Recent demographic history inferred by high-resolution analysis of linkage disequilibrium. Molecular Biology and Evolution, 37(12), 3642–3653.

Shahar, D.J., 2017. Minimizing the variance of a weighted average. Open Journal of Statistics, 7(2), pp.216–224.

Tenesa, A., Navarro, P., Hayes, B. J., Duffy, D. L., Clarke, G. M., Goddard, M. E., & Visscher, P.M. (2007). Recent human effective population size estimated from linkage disequilibrium. Genome Research, 17, 520–526.

Wang, J., 2001. A pseudo-likelihood method for estimating effective population size from temporally spaced samples. Genetics Research, 78(3), pp.243–257.

Wang J (2009) A new method for estimating effective population size from a single sample of multilocus genotypes. Molecular Ecology, 18, 2148–2164.

Wang, J., 2016. A comparison of single-sample estimators of effective population sizes from genetic marker data. Molecular Ecology, 25(19), pp.4692–4711.

Wang, J. and Santure, A.W., 2009. Parentage and sibship inference from multilocus genotype data under polygamy. Genetics, 181(4), pp.1579–1594.

Wang, J., E. Santiago, and A. Caballero. 2016. “Prediction and Estimation of Effective Population Size.” Heredity 117: 193–206.

Waples, R.S. 1989. A generalized approach for estimating effective population size from temporal changes in allele frequency. Genetics 121:379–391.

Waples, R.S. 2005. Genetic estimates of contemporary effective population size: To what time periods do the estimates apply? Molecular Ecology 14:3335–3352.

Waples, R.S. 2006. A bias correction for estimates of effective population size based on linkage disequilibrium at unlinked gene loci. Conservation Genetics 7:167–184.

Waples, RS. 2021. Relative precision of the sibship and LD methods for estimating effective population size with genomics-scale datasets. Journal of Heredity 112:535:539.

Waples, R.S. 2024. Practical application of the linkage disequilibrium method for estimating contemporary effective population size: a review. Molecular Ecology Resources 24:e13879

Waples, R.S. 2025. The Idiot’s Guide to effective population size. Molecular Ecology 10.1111/mec.17670

Waples, R.S., and C. Do. 2008. LdNe: A program for estimating effective population size from data on linkage disequilibrium. Mol. Ecol. Resources 8:753–756.

Waples, R.S., and C. Do. 2010. Linkage disequilibrium estimates of contemporary N_e_ using highly variable genetic markers: A largely untapped resource for applied conservation and evolution. Evolutionary Applications 3:244–262.

Waples, R.S., and J.R. Faulkner. 2009. Modeling evolutionary processes in small populations: Not as ideal as you think. Molecular Ecology 18:1834–1847.

Waples, R.S., Waples, R.K., and Ward, E.J., 2022. Pseudoreplication in genomics-scale datasets. Molecular Ecology Resources 22:503–518.

Waples, R.S., M.M. Masuda, M.E.F. LaCava, A.J. Finger. 2025. MaxTemp: A method to maximize precision of the temporal method for estimating N_e_ in genetic monitoring programs. Molecular Ecology Resources 25:e14057.

Weir, B.S., 1979. Inferences about linkage disequilibrium. Biometrics, pp.235–254.

Weir, B. S., & Hill, W. G. (1980). Effect of mating structure on variation in linkage disequilibrium. Genetics, 95(2), 477–488.

Wright, S., 1965. The interpretation of population structure by F-statistics with special regard to systems of mating. Evolution, pp. 395–420.

