## Supplemental tables and figs for "Combining multiple genetic estimates of *N*_*e*_"

Table S1. Comparison of var($1/\widehat{N}_e$) for two scenarios using the temporal method. Scenario 1: $\widehat{N}_e$ is computed using two samples taken *t* generations apart (as in Equation 11); Scenario 2: $\widehat{N}_{eCombo}$ is computed from two separate estimates, each based on two samples taken *t*/2 generations apart (as in Equation 24, assuming the covariance term = 0). Values shown in the table are the ratio of var($1/\widehat{N}_e$) for Scenario 2 to that of Scenario 1. Bolded values indicate parameter combinations for which Scenario 2 is expected to have larger variance, even assuming independence of the nested estimates. See Figure S3 for simulation results.

| | | Sample size (*S*) | | | |
|---|---|---|---|---|---|
| | $N_e$ | 25 | 50 | 100 | 200 |
| *t*=1 | 50 | **1.62** | **1.39** | **1.13** | 0.89 |
| | 100 | **1.78** | **1.62** | **1.39** | **1.13** |
| | 200 | **1.88** | **1.78** | **1.62** | **1.39** |
| | 500 | **1.95** | **1.91** | **1.82** | **1.68** |
| | 1000 | **1.98** | **1.95** | **1.91** | **1.82** |
| | 5000 | **2.00** | **1.99** | **1.98** | **1.96** |
| *t*=5 | 50 | **1.04** | 0.83 | 0.68 | 0.60 |
| | 100 | **1.30** | **1.04** | 0.83 | 0.68 |
| | 200 | **1.55** | **1.30** | **1.04** | 0.83 |
| | 500 | **1.78** | **1.62** | **1.39** | **1.13** |
| | 1000 | **1.88** | **1.78** | **1.62** | **1.39** |
| | 5000 | **1.98** | **1.95** | **1.91** | **1.82** |
| *t*=10 | 50 | 0.83 | 0.68 | 0.60 | 0.55 |
| | 100 | **1.04** | 0.83 | 0.68 | 0.60 |
| | 200 | **1.30** | **1.04** | 0.83 | 0.68 |
| | 500 | **1.62** | **1.39** | **1.13** | 0.89 |
| | 1000 | **1.78** | **1.62** | **1.39** | **1.13** |
| | 5000 | **1.95** | **1.91** | **1.82** | **1.68** |

Table S2. Confidence interval (CI) coverage for the example shown in Figure 4. The "Covered" column shows the fraction of 5000 simulated confidence intervals for combined estimates of $N_e$ that included the true value (1000). The "Low" and "High" columns show the fractions of CIs that were entirely too low or too high, respectively. r is the correlation between the two estimators, A and B. For r=0.5, CIs were computed two ways: x—based on theoretical variance using the full Equation 2; y—ignoring the covariance term in Equation 2.

| *r* | Low | Covered | High |
|---|---|---|---|
| 0 | 0.026 | 0.950 | 0.024 |
| 0.5x | 0.026 | 0.948 | 0.026 |
| 0.5y | 0.052 | 0.897 | 0.051 |

Table S3. Confidence interval (CI) coverage for the LD analyses shown in Figure S3. The "Covered" column shows the fraction of simulated confidence intervals for estimates of $N_e$ that included the true value (200). The "Low" and "High" columns show the fractions of CIs that were entirely too low or too high, respectively. *n* is the number of generations of data used in the estimate of $N_e$.

| *n* | Low | Covered | High |
|---|---|---|---|
| 1 | 0.022 | 0.938 | 0.040 |
| 2 | 0.017 | 0.920 | 0.064 |
| 5 | 0.011 | 0.874 | 0.116 |
| 10 | 0.001 | 0.807 | 0.192 |

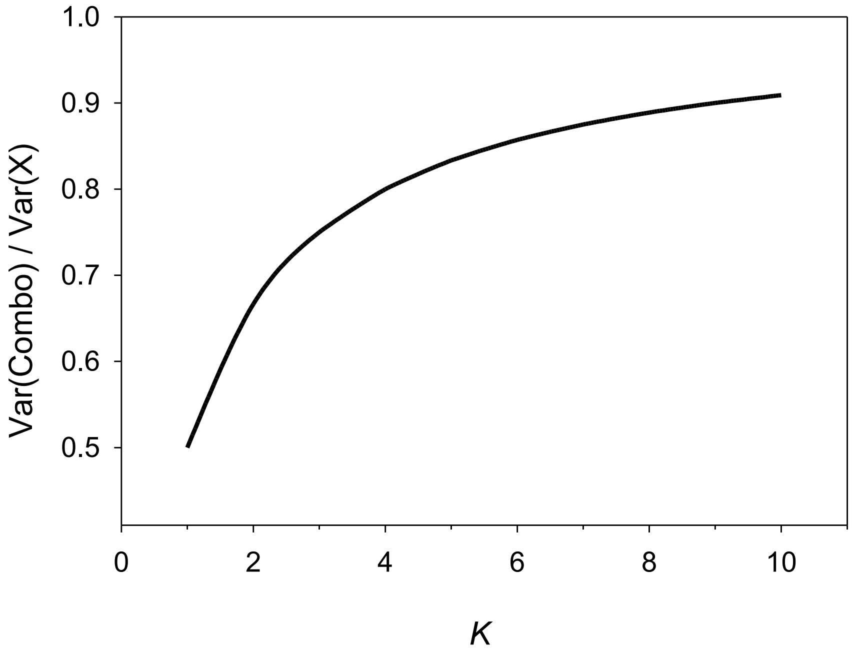


Figure S1. Variance of a combined estimator (Var(Combo)) as a proportion of the variance of the more precise of two individual estimators (estimator *X*). It is assumed that the estimators are uncorrelated and the variance of the less-precise estimator is *var*(*Y*) = *K*var(X)*, where *K*≥1. The curve depicts the relationship shown in Equation 28. Maximum reduction in variance (leading to maximum increase in precision) occurs when *K*=1 (so *var*(*Y*)=*var(X)*).

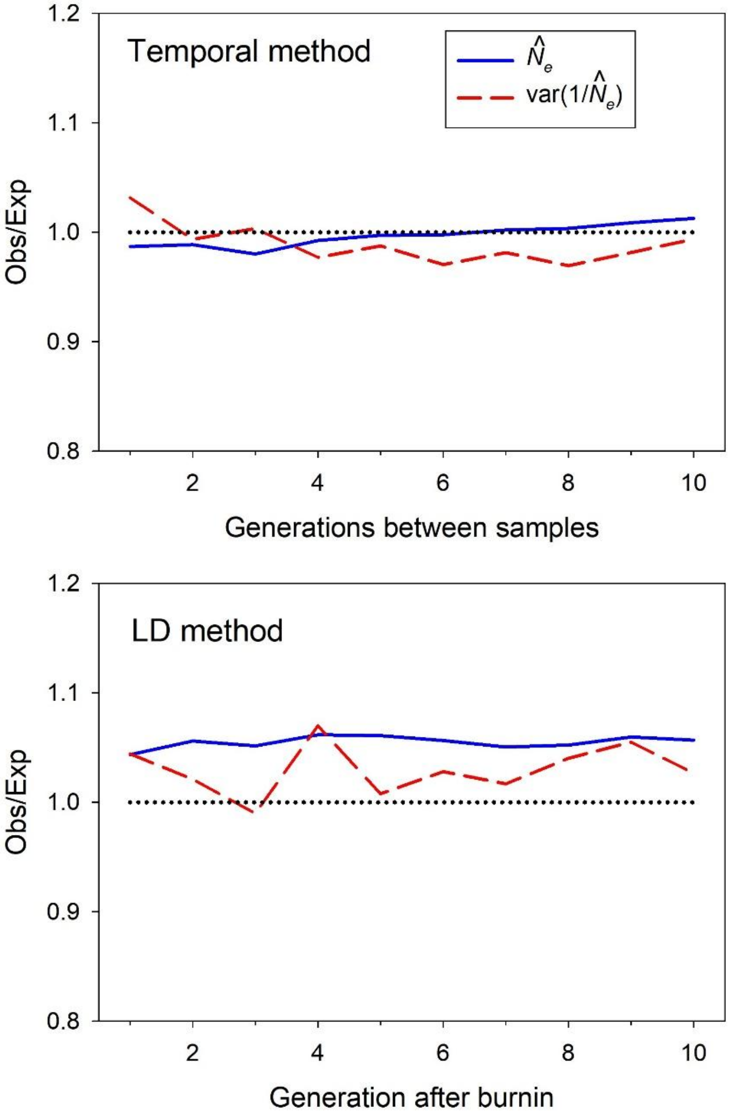


Figure S2. Within-method simulation results for the temporal method (top panel) and LD method (bottom panel). For both methods, each of 500 replicate simulations was run for 10 generations. True $N_e$ was 200, and data were computed across 20 replicate samples of 50 individuals at each generation, assayed for variation at 1000 diallelic loci. The Y axes show the ratio of observed to expected values for harmonic mean $\widehat{N}_e$ (solid blue lines) and variance of $1/\widehat{N}_e$ (dashed red lines). For the temporal method, results are averaged across all pairwise comparisons off samples taken 1 ... 10 generations apart, and expected values of var($1/\widehat{N}_e$) were computed from Equation 11. For the LD method, results are averaged across samples taken 1 ... 10 generations after the burnin, all of which have the same expected value of var($1/\widehat{N}_e$) given by Equation 16.

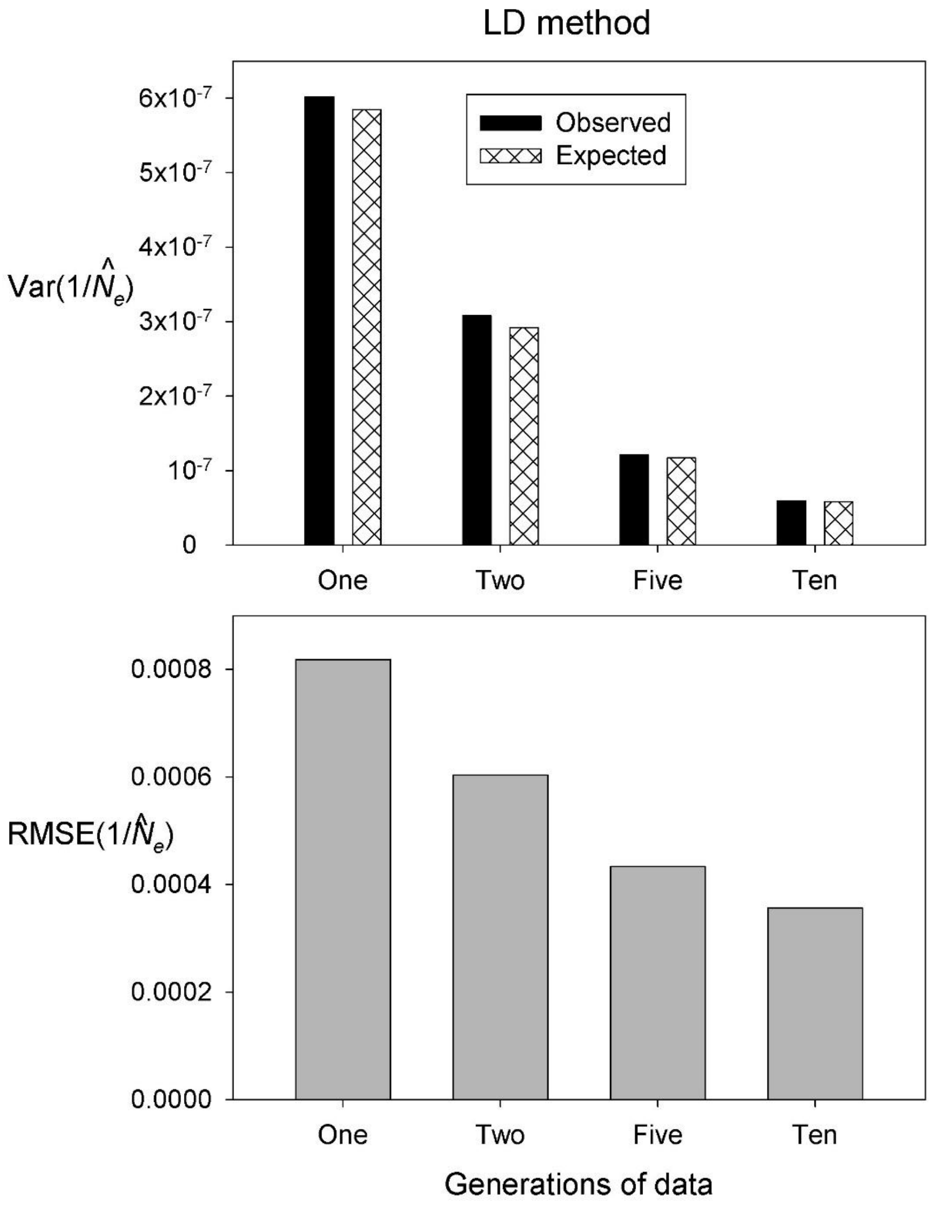


Figure S3. Top: Comparison of observed and expected values of var($1/\widehat{N}_e$) for the LD method when one, two, five, or ten generations of samples are used to compute an overall $\widehat{N}_e$. As in Figure S2, true $N_e$ was 200, and data were computed across 20 replicate samples of 50 individuals at each generation, assayed for variation at 1000 diallelic loci. Observed values were computed from simulations, and expected values were computed using Equation 23 assuming no covariance across generations. Bottom: RMSE of $1/\widehat{N}_e$ for individual estimates or combo estimates using 2, 5, or 10 generations of data.

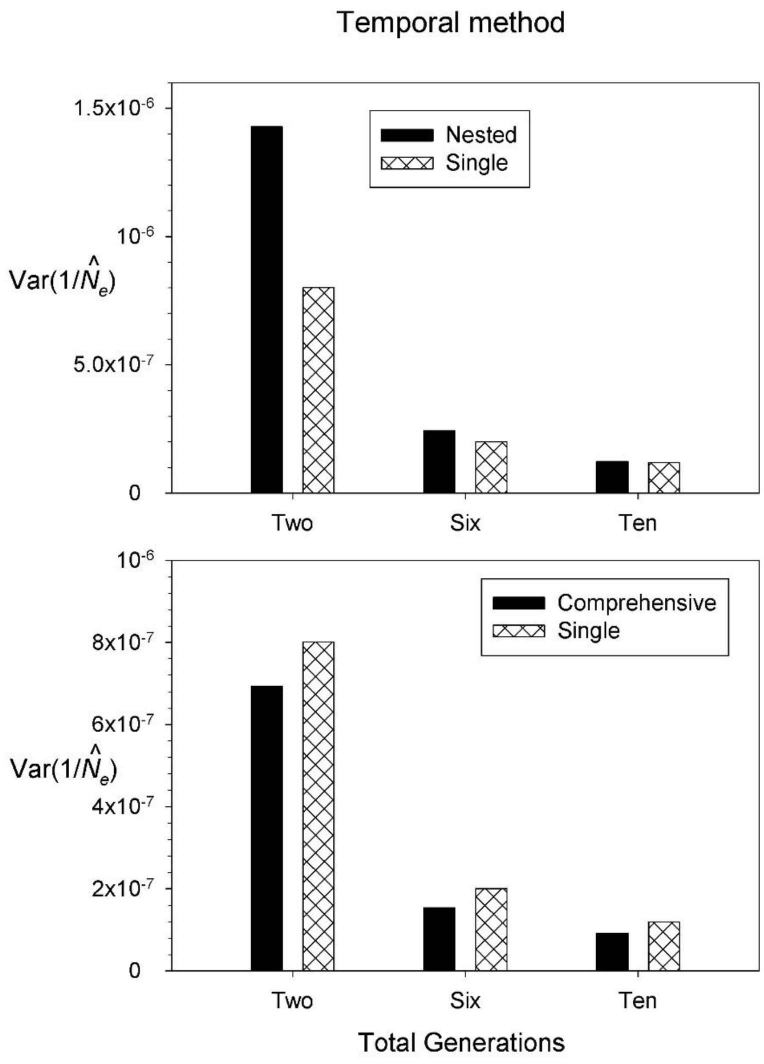


Figure S4. Comparison of observed and expected values of var($1/\widehat{N}_e$) for the temporal method when results from multiple estimates are combined. Simulation parameters were as in Figure S2. In both panels, the reference vertical bars (cross-hatched) show results for a single pair of samples spanning the indicated number of generations (2, 5, or 10). In the top panel, the black vertical bars show results for nested, combined estimates. For two generations, the nested results use equally-weighted data for comparisons of generations 0 and 1 and generations 1 and 2; for ten generations, the nested results use equally-weighted data for comparisons of generations 0 and 5 and generations 5 and 10; for five generations, the nested results use comparisons of generations 0 and 2 and generations 2 and 5, weighted as in Equation 24. In the bottom panel, the black bars show results when the nested estimates are combined with the single estimates to produce a comprehensive estimate that uses data for all of the comparisons.
